# Comments on a small sabretooth cat in the Abismo Ponta de Flecha Cave, Vale do Ribeira, southeastern Brazil

**DOI:** 10.1101/2021.09.03.458923

**Authors:** Artur Chahud

**Affiliations:** Laboratory for Human Evolutionary Studies, Department of Genetics and Evolutionary Biology, Institute of Biosciences, University of São Paulo. Rua do Matão 277, São Paulo, SP 05508-090, Brazil.

**Keywords:** Felidae, Pleistocene, Carnívora, South America, Machairodontinae

## Abstract

The Vale do Ribeira, located in southeastern Brazil, is known for many caves with osteological material, including several extinct species. The saber-tooth cat *Smilodon populator* was a large predator that inhabited the Pleistocene and Holocene of South America. A specimen found in the Abismo Ponta de Flecha Cave based on small bones (metacarpals and phalanges) is commented here. The metacarpals have morphological characteristics of *S. populator*, but are smaller than that of *S. fatalis* and *Panthera onca* and larger than that of *S. gracilis*. The specimen is among the smallest ever found and is comparable in size to an adult lion.

## 1. Introduction

The Vale do Ribeira, located in southeastern Brazil, gathers carbonate rocks where a complex system of caves developed. Despite the high paleontological potential of this karst region, a few works involving the study of fossils have been developed there.

The Abismo Ponta de Flecha is a vertical cave located in ancient centripetal drainage, formed in carbonate rocks of the Proterozoic (Barros Barreto *et al*., 1982). The site has a large amount of osteological material, composed of extinct and living animals (Barros-Barreto *et al*., 1982; Chahud, 2005).

The Felidae occurs in South America since the Great American Biotic Interchange 2 (GABI 2), which occurred on the boundary between the Pliocene and Pleistocene, 1.8 Ma (Woodburne, 2010), with the migration of several species from North America.

Currently, the large Felidae from South America are represented by only two species, *Puma concolor* and *Panthera onca*. However, several other species are represented, all coming from the Late Pleistocene and Early Holocene.

Among the extinct genera reported in South America, the most common is the subfamily Machairodontinae, represented by the genera; *Smilodon, Homotherium* and *Xenosmilus*, being the first most common and the only one with representatives found in Brazil, while the others are local occurrences in Venezuela and Uruguay, respectively (Mones & Rinderknecht, 2004; RincÓn *et al*., 2011).

For a long time, *Smilodon populator* was the only representative of the genus in South America, but evidence of other species has been reported; *S. fatalis* has been found in the Andean region of Ecuador, Peru and Uruguay (KurtÉn & Werdelin, 1990; Manzuetti *et al*., 2018), while *S. gracilis* has been described in the Venezuelan Andes (Rincón *et al*., 2011).

The species *Smilodon populator* was originally described by Lund (1842) of specimens described in Lagoa Santa region, where several other specimens were found in several caves (Hubbe *et al*., 2013; Chahud, 2020). In the Vale do Ribeira region, other specimens have been described, and the specimen found in the Abismo Iguatemi Cave presented the best specimen (Ferreira & Karmann, 2002; Castro & Langer, 2008; 2011), revealing important information about the species.

The presence of an indeterminate Felidae was first noted by Chahud (2005), based on bone materials initially identified as Xenarthra. The present work identifies this specimen as *Smilodon populator* and makes comparisons with others described in the Quaternary of South America.

## 2. Material and Methods

The osteological material of the Abismo Ponta de Flecha Cave consists of more than 1400 samples, including faunal and inorganic remains, and was collected by a team of geologists and biologists between 1981-1982, as part of a large speleological study, archaeological to paleontological in Vale do Ribeira (Figure 1). Much of this material is cured, including the specimens studied here, in the Laboratory of Systematic Paleontology of the Department of Sedimentary and Environmental Geology of the Geosciences Institute – USP.

**Figure 1.**
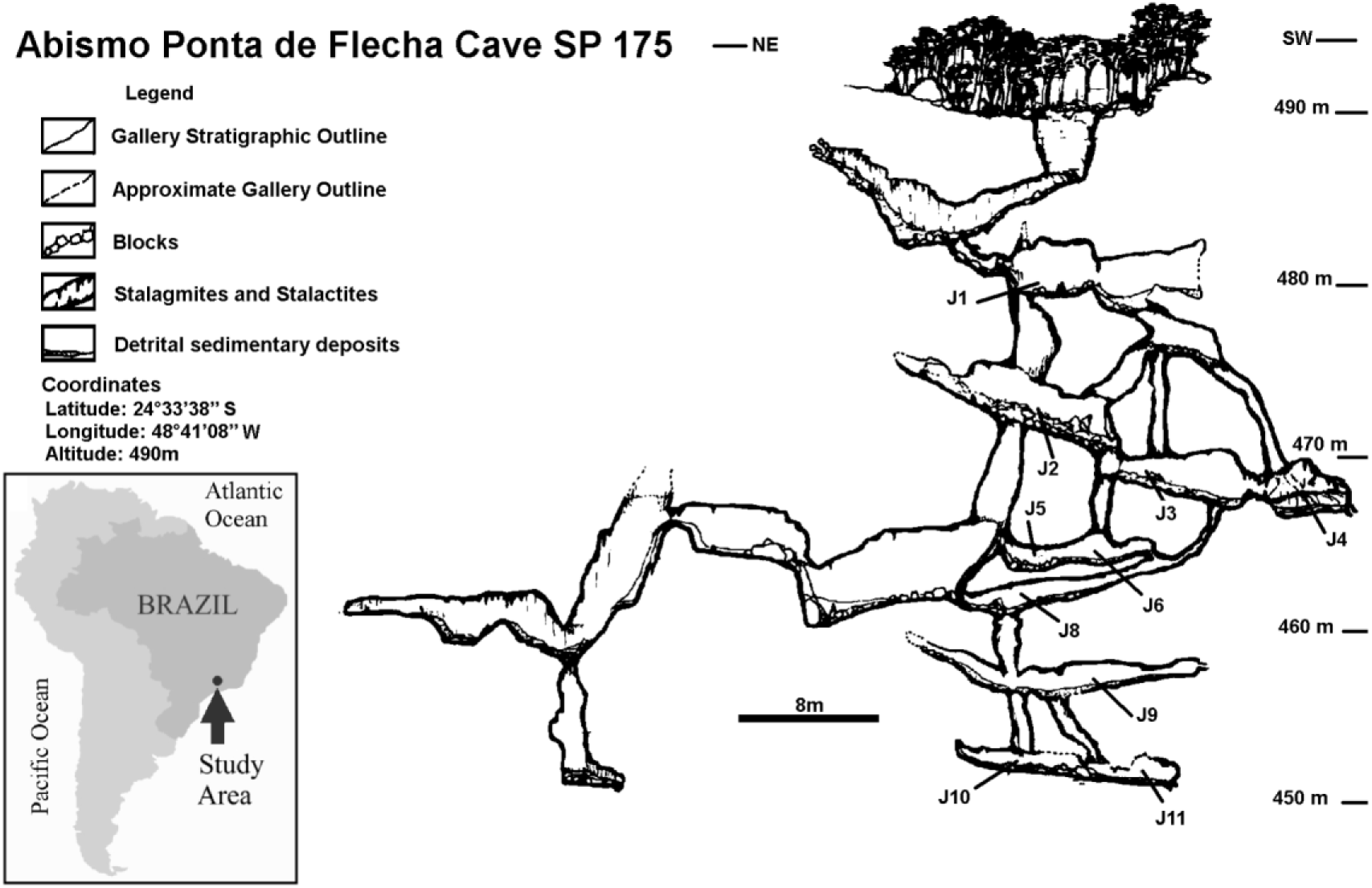
Schematic profile of the Abismo Ponta de Flecha, SP 175. Highlighting the galleries (*Jazidas*) with osteological material J1-J11, (Adapted from Barros-Barreto *et al*., 1982).

Initially, the specimens were organized and listed according to location and positioning in the gallery, called *Jazidas*, in which each piece was found (PF-), later the material received a second registration number (GP/2C-). The study specimens come from *Jazida* 2 (J2), characterized by ancient sedimentation levels and collapsed blocks (Figure 1). Below these blocks, fossil bones of large animals were found (Barros Barreto *et al*., 1982).

## 3. Systematic Palaeontology

**Order Carnivora Bowdich, 1821**

**Family Felidae Gray, 1821**

**Subfamily Machairodontinae Gill, 1872**

**Tribe Smilodontini Kurtén, 1963**

**Genus *Smilodon* Lund, 1842**

**Figure 2, Figure 3**

**Figure 2.**
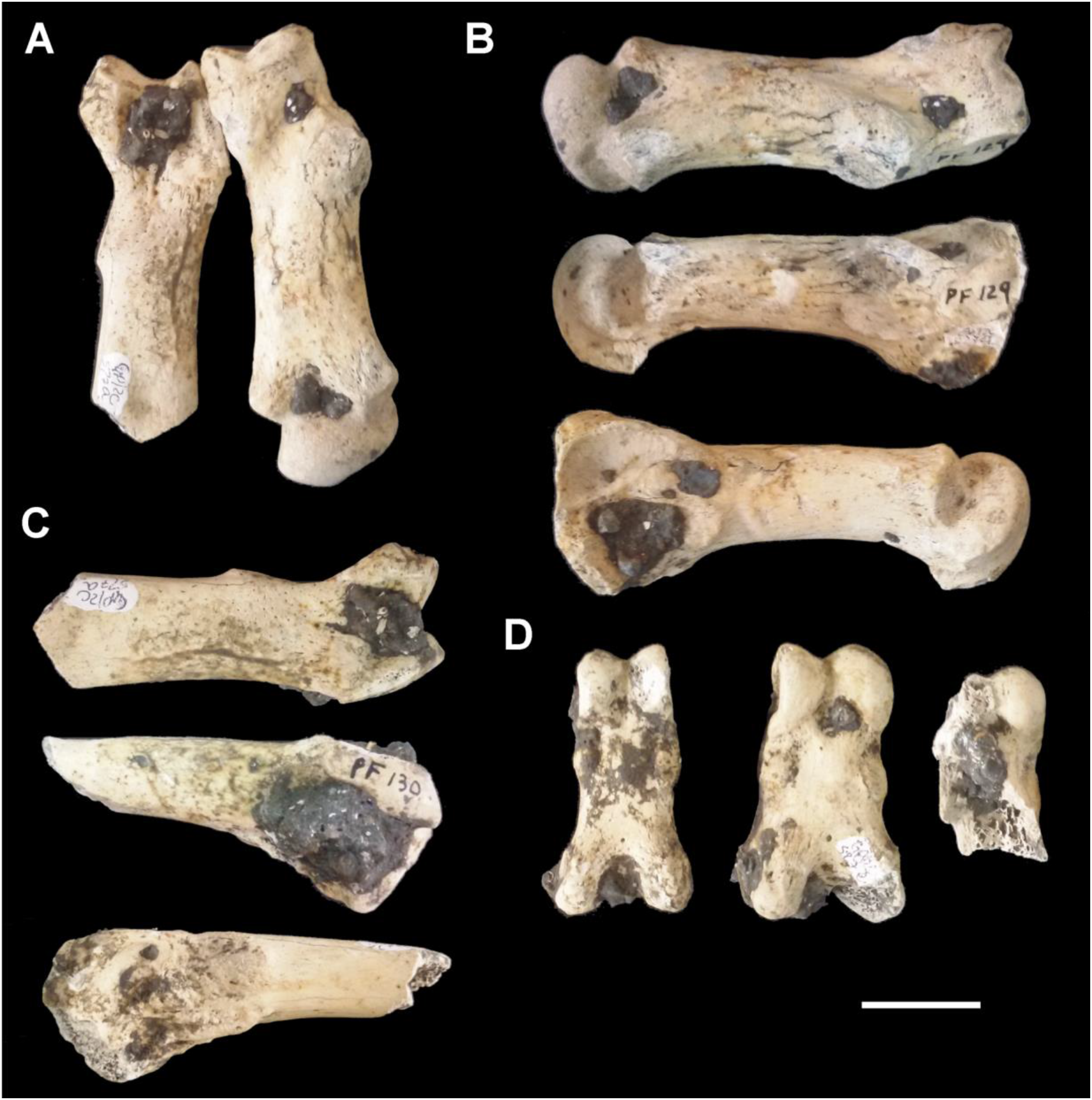
*Smilodon populator* from the Abismo Ponta de Flecha Cave. A) Metacarpals III and II (PF 130–GP/2C-527a, PF 129–GP/2C-527d), dorsal view; B) Metacarpals II dorsal and lateral views. C) Metacarpal III dorsal and lateral views. D) Palmar view of the proximal phalanges II, III and IV (PF 132 – GP/2C-527b, PF 131 – GP/2C-527c, PF 133 – GP/2C-527e). Scale: 20mm.

**Figure 3.**
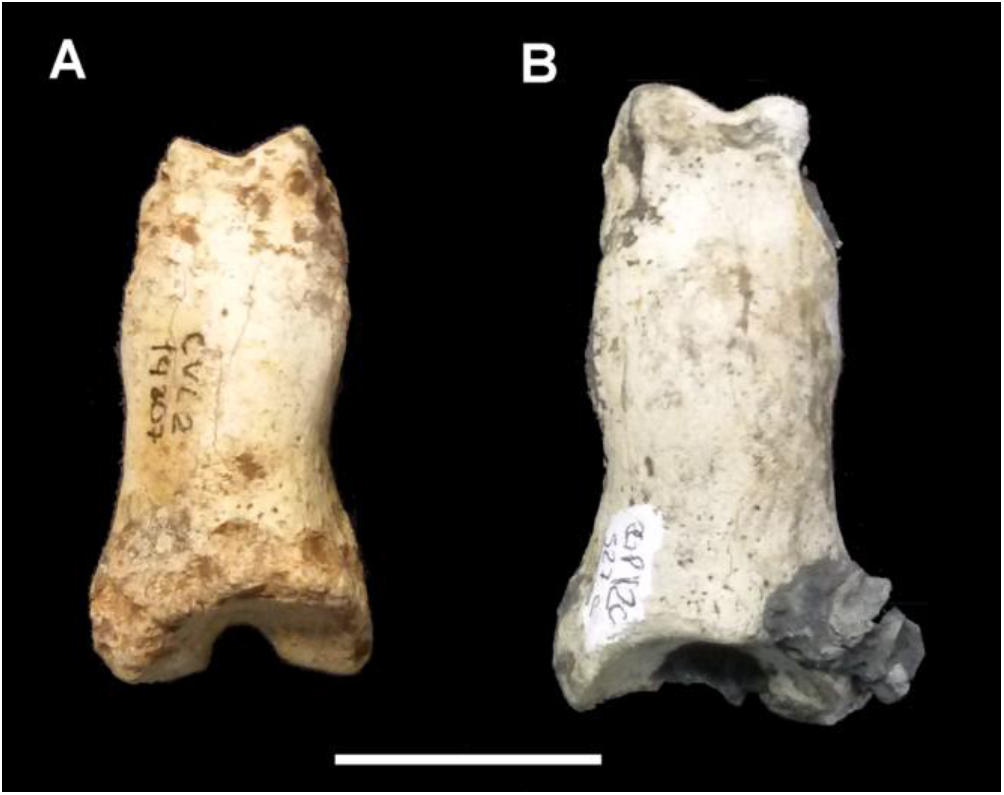
Dorsal views of proximal phalanges of *Smilodon populator*. A) Cuvieri Cave (CVL2-14207); B) Abismo Ponta de Flecha Cave. Scale: 20mm.

### Type species

*Smilodon populator* Lund, 1842

### Material

The identified material consists of a complete left metacarpal II (PF 129–GP/2C-527d) (Figure 2A and 2B), fragmented left metacarpal III (PF 130–GP/2C-527a) (Figure 2A and 2C), two complete proximal left phalanges (PF 131–GP/2C-527c, PF 133–GP/2C-527e) and fragmented proximal left phalanx (PF 132–GP/2C-527b) (Figure 2D).

### Taphonomy

The osteological material assigned to the specimen is whitish and with carbonate incrustation (Figure 2A – C), which could not be removed without compromising the structure of the bone part. No cracks caused by bone exposure or polishing were observed, suggesting that the specimen was not transported into the cave after its death and that its presence in its interior was probably accidental. The hypothesis is that the specimen entered the cave alive and became trapped inside it.

### General characteristics and comparison

#### Proximal Phalanges

The recovered osteological material is represented by the left proximal phalanges II, III and IV (Figure 2D). The proximal phalanges II and III have sizes and shapes compatible with phalanges attributed to large cats that inhabited South America during the Pleistocene and compared to a proximal phalanx, but of the hind limb, of a specimen from Cuvieri cave, eastern Brazil (Figure 3), the specimen is larger, but compared to the phalanges described by Méndez-Alzola (1941), a *Smilodon* found in the Argentine Pampa, the specimen is smaller (Table 1). Phalanx IV was very fragmented and no measurements or comparisons were possible.

**Table 1.** Measures of proximal phalanges II and III in specimens found in the Abismo Ponta de Flecha Cave, Cuvieri Cave and Argentine Pampa

|  | Abismo Ponta de Flecha |  | Cuvieri Cave | Argentine Pampa (Méndez-Alzola, 1941) |  |
| --- | --- | --- | --- | --- | --- |
|  | II | III | II | II | III |
| Length | 46.3 | 44.95 | 40.2 | 52 | 55.8 |
| Shaft width | 18,5 | 19,9 | 16.08 | 24.9 | 18 |
| Shaft deep | 11.54 | - | 11.67 | 15.5 | 14.3 |
| Distal width | 15.7 | 17.81 | 15.8 | 20.2 | 20 |
| Distal Deep | 11.23 | 13.4 | 10.41 | - | - |
| Proximal width | 23.18 | 23.6 | 19.93 | 27.6 | 28.3 |
| Proximal deep | 17.53 | 17.5 | 17.04 | 23.3 | 21 |

#### Metacarpals

The metacarpal II of the specimen found in the Abismo Ponta de Flecha Cave is more robust but smaller than the metacarpals of Pleistocene and recent species of the genus *Panthera* (Chahud & Okumura, 2020). Compared with the *Smilodon* specimens studied by KurtÉn & Werdelin (1990) (Table 2) it is larger and more robust than *S. gracilis*. The width is comparable to specimens of *S. fatalis*, both from North and South America, but the length is smaller than individuals of this species, according to KurtÉn & Werdelin (1990) this characteristic is expected in specimens of *S. populator*.

**Table 2.** Measures of metacarpals bones of the Abismo Ponta de Flecha Cave specimen and other *Smilodon* species studied by KurtÉn & Werdelin (1990).

|  | *Smilodon*<br>Abismo<br>Ponta de<br>Flecha | *Smilodon*<br>*fatalis*<br>Rancho<br>La brean | *Smilodon*<br>*fatalis*<br>Talara,<br>Peru | *Smilodon*<br>*populator*<br>Brazil and<br>Argentina | *Smilodon*<br>*gracilis* |
| --- | --- | --- | --- | --- | --- |
| **Metacarpal II** | N = 1 | N=7 | N = 19 | N= 4 | N = 3 |
| Length | 76.6 | 90.6 ± 4.4 | 85.5 ± 1.1 | 89.2 ± 1.4 | 73.6 ± 1.3 |
| Shaft width | 16.97 | 17.2 ± 0.5 | 16.5 ± 0.3 | 19.4 ± 0.4 | 12.4 ± 0.5 |
| Shaft deep | 15.53 |  |  |  |  |
| Distal width | 25.2 | 24.8 ± 0.8 | 24.3 ± 0.3 | 26.6 ± 0.2 | 19.2 ± 0.9 |
| Distal Deep | 22.82 |  |  |  |  |
| Proximal width | 25.27 |  |  |  |  |
| Proximal deep | 30.31 |  |  |  |  |
| **Metacarpal III** | N = 1 | N=7 | N = 19 | N= 3 | N = 1 |
| Length |  | 96.4 ± 4.0 | 97.3 ± 0.9 | 97.3 ± 2.7 | 73 |
| Shaft width | 16.2 | 17.3 ± 0.8 | 16.4 ± 0.3 | 19.4 ± 0.7 | 10.5 |
| Shaft deep | 12.23 |  |  |  |  |
| Distal width |  | 26.5 ± 1.0 | 25.6 ± 0.4 | 28.2 ± 0.2 | 16.7 |
| Proximal width | 25.8 |  |  |  |  |
| Proximal deep | 26.41 |  |  |  |  |

Compared with specimens of *Smilodon populator* from Argentina, the specimen from Vale do Ribeira is morphologically similar to *S. populator* described by KurtÉn & Werdelin (1990), but smaller, being probably a small individual of this species. The relationship between shaft width and length of *Smilodon populator* specimens of KurtÉn & Werdelin (1990) is 0.21749 while that of Abismo Ponta de Flecha Cave is 0.22154, which are similar values when compared to other *Smilodon* species (Table 3), it can be associated with the species *S. populator*.

**TABLE 3.**
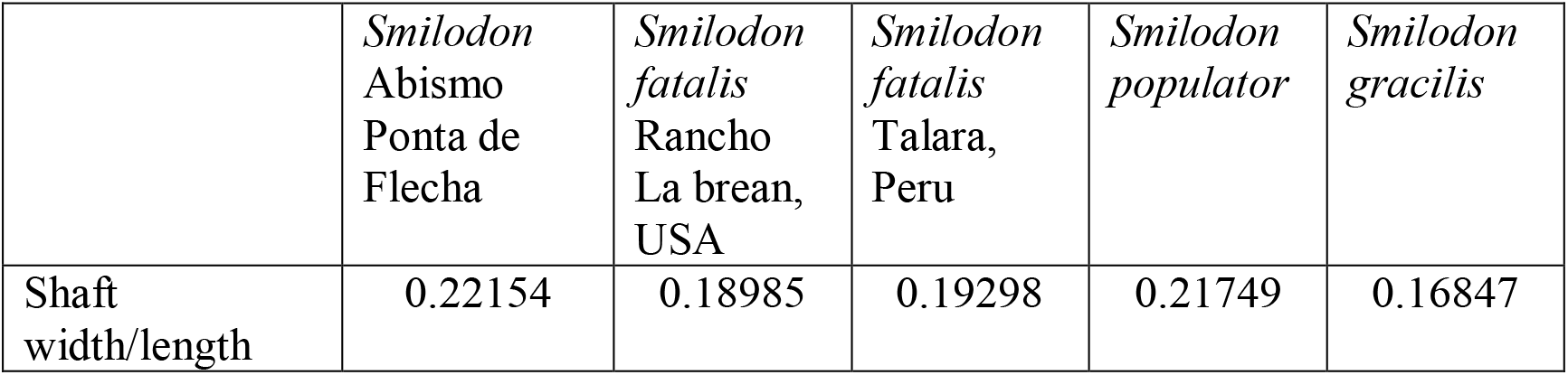
Relationship between the shaft width and the length of *Smilodon* species by KurtÉn & Werdelin (1990) compared to the specimen found in Abismo Ponta de Flecha Cave.

|  | *Smilodon*<br>Abismo<br>Ponta de<br>Flecha | *Smilodon*<br>*fatalis*<br>Rancho<br>La brean,<br>USA | *Smilodon*<br>*fatalis*<br>Talara,<br>Peru | *Smilodon*<br>*populator* | *Smilodon*<br>*gracilis* |
| --- | --- | --- | --- | --- | --- |
| Shaft<br>width/length | 0.22154 | 0.18985 | 0.19298 | 0.21749 | 0.16847 |

## 4. Discussion and Final Considerations

The specimen recovered from the Abismo Ponta de Flecha Cave represents a *Smilodon* of small proportions, compared to other specimens from South America.

The genus *Smilodon* reported differences in proportion and morphology between the three species, and regional morphological differences in *S. fatalis* (Tables 2 and 3) from North America and Peru (KurtÉn & Werdelin, 1990), and the same may have occurred with *S. populator*.

Several mammals vary in size between individuals of the same species or genus. In Africa, a variation in the proportion between the two species of elephants of the genus *Loxodonta* has been reported, with *L. cyclotis*, which inhabits forested regions, is smaller than *L. africana* (Grubb *et al*., 2000) found in savannas. The same was reported for South American species of the genus *Tapirus*, among *T. terrestris* (more common), *T. pinchaque* (from the Andean region and smaller) and *T. bairdii* (from Central America and larger) (Ruiz-GarcÍa *et al*. 2016).

In South America, size variations between individuals of the same species have been reported, as observed in *Dicotyles tajacu* from the Amazon region, compared to other regions (Gongora *et al*. 2011). However, among Felidae, the greatest difference was observed in small individuals of the species *Puma concolor* from the Amazon, compared to specimens from the Andean region and southern South America (Pacheco & Zapata, 2017; Chimento & Dondas, 2018; Chahud, 2021).

The species *Smilodon populator* is the largest of the genus and larger than the largest recent felines, but it is possible to find individuals with an approximate size of *S. fatalis*, while the largest lions (*Panthera leo*) are proportionally comparable to the smallest *S. populator* (Christiansen & Harris, 2005). The specimens found in the Cuvieri Cave (Chahud, 2020) and in the Abismo Ponta de Flecha Cave are of comparable size to *S. fatalis* and recent lions and are probably among the smallest *S. populator*.

## Acknowledgements

The author thanks M.A. Aragão for support. The author also thanks Dr. M. Mercedes Martinez Okumura responsible for LEEH (Laboratory for Human Evolutionary Studies), Department of Genetics and Evolutionary Biology, Institute of Biosciences of the University of São Paulo which allowed the preparation of fossils in her laboratory. AC holds a CNPq Senior Post-doctoral scholarship (103934/2020-0)

## Bibliographic References

Barros Barreto, C.N.G., de Blasiis, P.A.D., Dias Neto, C.M., Karmann, I., Lino, C.F. & Robrahn, E.M., 1982. Abismo Ponta de Flecha: um projeto arqueológico, paleontológico e geológico no médio Ribeira de Iguape, SP. Revista da Pré História, vol. 3, p. 195–215.

Castro, M.C. & Langer, M. C., 2008. New postcranial remains of *Smilodon populator* Lund, 1842 from southeastern Brazil. Revista Brasileira de Paleontologia, vol. 11. p. 199–206. https://doi.org/10.4072/rbp.2008.3.06

Castro, M.C. & Langer, M.C., 2011. The mammalian fauna of Abismo Iguatemi, southeastern Brazil. Journal of Cave and Karst Studies, vol. 73, n. 83-92. https://doi.org/10.4311/jcks2010pa0140

Chahud, A. 2005. Paleomastozoologia do Abismo Ponta de Flecha, Iporanga, SP. In: II Congresso Latino-Americano de Paleontologia de Vertebrados. Abstracts. Rio de Janeiro: Museu Nacional/UFRJ, p. 76–78.

Chahud, A. 2020. Occurrence of the sabretooth cat *Smilodon populator* (Felidae, Machairodontinae) in the Cuvieri cave, eastern Brazil. Palaeontologia Electronica, vol. 23, n. 2: a24. https://doi.org/10.26879/1056

Chahud, A. 2021. Osteometria e breves comentários sobre *Puma concolor* Linnaeus, 1771 (Carnivora, Felidae), no Estado do Maranhão, Brasil. Revista Biociências - Universidade de Taubaté, vol. 27 n. 1, p. 15–28.

Chahud, A. & Okumura, M. 2020. The presence of *Panthera onca* Linnaeus 1758 (Felidae) in the Pleistocene of the region of Lagoa Santa, State of Minas Gerais, Brazil. Historical Biology. https://doi.org/10.1080/08912963.2020.1808975

Chimento, N. R. & Dondas, A. 2018. First record of *Puma concolor* (Mammalia, Felidae) in the Early-Middle Pleistocene of South America. Journal of Mammalian Evolution, vol. 25, n. 3, p. 381–389. https://doi.org/10.1007/s10914-017-9385-x

Christiansen, P. & Harris, J. M. 2005. Body size of *Smilodon* (Mammalia: Felidae). Journal of Morphology, vol. 266, n. 3, p. 369–84. https://doi.org/10.1002/jmor.10384

Ferreira, N.B. & Karmann, I. 2002. Descobertas paleontológicas na região de Apiaí-SP. Boletim Informativo Geovisão, vol. 10, n. 4, p. 7–8.

Gongora, J., Biondo, C., Cooper, J.D., Taber, A., Keuroghlian, A., Altrichter, M., Ferreira do Nascimento, F., Chong, A.Y., Miyaki, C.Y., Bodmer, R., Mayor, P. & González, S. 2011. Revisiting the species status of *Pecari maximus* van Roosmalen et al., 2007 (Mammalia) from the Brazilian Amazon. Bonn Zoological Bulletin, vol. 60, n. 1, p. 95–101.

Grubb, P.; Groves, C. P.; Dudley, J. P. & Shoshani, J. 2000. Living African elephants belong to two species: *Loxodonta africana* (Blumenbach, 1797) and *Loxodonta cyclotis* (Matschie, 1900). Elephant, vol. 2, n. 4, p. 1–4. doi:10.22237/elephant/1521732169

Hubbe, A., Hubbe, M., Karmann, I., Cruz, F.W., & Neves, W.A. 2013. Insights into Holocene megafauna survival and extinction in southeastern Brazil from new AMS 14C dates. Quaternary Research, vol. 79, p. 152–157. https://doi.org/10.1016/j.yqres.2012.11.009

Kurtén, B. & Werdelin, L. 1990. Relationships between North and South American Smilodon. Journal of Vertebrate Paleontology, vol. 10, n. 2, p. 158–169. https://doi.org/10.1080/02724634.1990.10011804

Lund, P.W. 1842. Blik paa Brasiliens Dyreverden for Sidste Jordomvaeltning. Tredie Afhandling: Forsaettelse af Pattedyrene. Det Kongelige Danske Videnskabernes Selskabs Naturvidenskabelige og Mathematiske Afhandlinger, vol. 9, p. 137–208.

Manzuetti A., Perea D., Ubilla M., & Rinderknecht A., 2018. First record of *Smilodon fatalis* Leidy, 1868 (Felidae, Machairodontinae) in the extra-Andean region of South America (late Pleistocene, Sopas Formation), Uruguay: taxonomic and paleobiogeographic implications. Quaternary Science Reviews, vol. 180, n. 57-62. https://doi.org/10.1016/j.quascirev.2017.11.024

Méndez-Alzola, R. 1941. El *Smilodon bonaërensis* (Muñiz), Estudio osteológico y osteométrico del gran tigre fósil de La pampa comparado con otros félidos actuales y fósiles. Anales del Museo Argentino de Ciencias Naturales “Bernardino Rivadavia”, vol. 40, n. 67, p. 135–252.

Mones, A. & Rinderknecht, A. 2004. The First South American Homotheriini (Mammalia: Carnivora: Felidae). Comunicaciones Paleontologicas Museo Nacional de Historia Natural y Anthropologia, vol. 2, n. 35, p. 201–212.

Pacheco, J.I. & Zapata, C. 2017. Descripción osteológica del puma andino (Puma concolor): I. Esqueleto Apendicular. Revista de Investigaciones Veterinarias del Peru, vol. 28, n. 4, p. 1047–1054.

Rincón, A.D., Prevosti, F.J. & Parra, G.E. 2011. New saber-toothed cat records (Felidae: Machairodontinae) for the Pleistocene of Venezuela, and the Great American Biotic Interchange. Journal of Vertebrate Paleontology, vol. 31, n. 2, p. 468–478. https://doi.org/10.1080/02724634.2011.550366

Ruiz-García, M., Castellanos, A., Bernal, L.A., Pinedo-Castro, M., Kaston, F. & Shostell, J.M. 2016. Mitogenomics of the mountain tapir (*Tapirus pinchaque*, Tapiridae, Perissodactyla, Mammalia) in Colombia and Ecuador: Phylogeography and insights into the origin and systematics of the South American tapirs. Mammalian Biology, vol. 81, n. 2, p. 163–175. doi:10.1016/j.mambio.2015.11.001

Woodburne, M. O. 2010 The great American Biotic Interchange: Dispersals, Tectonics, Climate, Sea Level and Holding Pens. Journal of Mammalian Evolution, vol. 17, n. 4, p. 245–264.

